# Damage-induced pyroptosis drives endogenous thymic regeneration via induction of Foxn1 by purinergic receptor activation

**DOI:** 10.1101/2023.01.19.524800

**Authors:** Sinéad Kinsella, Cindy A. Evandy, Kirsten Cooper, Antonella Cardinale, Lorenzo Iovino, Paul deRoos, Kayla S. Hopwo, Colton W. Smith, David Granadier, Lucas B. Sullivan, Enrico Velardi, Jarrod A. Dudakov

## Abstract

Endogenous thymic regeneration is a crucial process that allows for the renewal of immune competence following stress, infection or cytoreductive conditioning. Fully understanding the molecular mechanisms driving regeneration will uncover therapeutic targets to enhance regeneration. We previously demonstrated that high levels of homeostatic apoptosis suppress regeneration and that a reduction in the presence of damage-induced apoptotic thymocytes facilitates regeneration. Here we identified that cell-specific metabolic remodeling after ionizing radiation steers thymocytes towards mitochondrial-driven pyroptotic cell death. We further identified that a key damage-associated molecular pattern (DAMP), ATP, stimulates the cell surface purinergic receptor P2Y2 on cortical thymic epithelial cells (cTECs) acutely after damage, enhancing expression of *Foxn1*, the critical thymic transcription factor. Targeting the P2Y2 receptor with the agonist UTPγS promotes rapid regeneration of the thymus *in vivo* following acute damage. Together these data demonstrate that intrinsic metabolic regulation of pyruvate processing is a critical process driving thymus repair and identifies the P2Y2 receptor as a novel molecular therapeutic target to enhance thymus regeneration.

**SUMMARY:**

- Thymocytes rapidly and transiently undergo pyroptosis after acute thymic damage and promote regeneration.
- Damage-induced redirection of pyruvate acutely enhances mitochondrial OXPHOS in thymocytes.
- Elevated mitochondrial ROS promotes pyroptosis in thymocytes after acute insult by driving caspase 1 cleavage.
- Extracellular ATP release promotes *Foxn1* expression in cTECs via activation of P2Y2
- Therapeutic targeting of the P2Y2 receptor promotes thymic regeneration.

## INTRODUCTION

Competent T cell development relies on efficient functioning of the thymus, which is extremely sensitive to acute insults, such as that caused by cytoreductive therapies^1^. Thymic function progressively declines with age, resulting in reduced export of newly generated naïve T cells and reduced responsiveness to new antigens and vaccines^2, 3^. The thymus has a remarkable ability to endogenously regenerate^4, 5, 6^, however, age-related deterioration drastically erodes this regenerative capacity^7^. Harnessing this regenerative capacity has the potential to expedite reconstitution of naïve T cells and improve immune responses. However, much remains unknown about the molecular regulators of this critical process.

We have previously identified that IL-22 and BMP4 represent two distinct pathways that facilitate endogenous repair in the thymus and, at least in the case of BMP4, is largely mediated by induction in the expression of Foxn1^5, 8^. FOXN1 is the essential thymic epithelial cell (TEC) transcription factor; not only crucial for the generation and function of TECs, but also for TEC maintenance with declining expression associated with age-related thymic involution^9^. We have previously identified that the constitutively high levels of homeostatic apoptosis in the steady state thymus, which governs negative selection events, is suppressive to the production of BMP4 and IL-23 (the upstream regulator of IL-22), and the depletion of apoptotic thymocytes after injury promotes their production^10^. Cell death is a sophisticated and tightly controlled process and much of this is regulated by the mitochondria, and notably necrotic cell death has been tightly linked to regeneration^11, 12, 13, 14, 15, 16^.

Given the robust depletion of thymocytes after acute damage concurrent to the activation of these reparative pathways, we hypothesized that a switch to an alternative cell death mechanism may underpin the triggering of tissue regeneration and alleviate the suppressive impact of apoptosis in the thymus. Here we investigated the effects of acute damage on the metabolic landscape of thymocytes and revealed that increased levels of pyruvate are redirected to mitochondrial respiration, reducing glycolysis. This disrupted glycolytic flux drives pyroptosis in the thymus which is rapidly resolved as regeneration begins.

These findings identify a novel mechanism of metabolic regulation of T cell development and thymic repair and provides a highly targetable therapeutic strategy to enhance immune function.

## RESULTS

### DP thymocytes preferentially undergo pyroptosis after damage

Most thymocytes, and in particular CD4+CD8+ double positive (DP) thymocytes, undergo apoptosis as a function of the selection processes fundamental to T cell development^17, 18^. We previously identified that homeostatic detection of these apoptotic events suppresses the production of multiple regenerative molecules in the thymus by promoting activation of TAM receptors bridging phosphatidylserine (PtdSer) sensing by surrounding stromal cells^10^. However, although we had previously found that after acute damage there is a rapid decrease in the detection of PtdSer^10^, this declined more rapidly than cell depletion itself (**Fig. 1A**) which led us to hypothesize that alternate forms of cell death may be being induced after acute damage. Given that DP thymocytes are the most numerous, comprising ∼80% of a thymus at baseline, and these cells are extremely sensitive to damage and are depleted rapidly after sub-lethal total body irradiation (TBI, 550 cGy) (**Fig. 1B**), we concentrated on this population. Not surprisingly we did find considerable cleavage of caspase-3 (executioner apoptosis caspase) within thymocytes (**Fig. 1C**), consistent with previous reports demonstrating their sensitivity to damage^19, 20^. However, we also found significant cleavage of caspase-1 in dying cells suggesting that in addition to immunologically silent apoptosis, there is also considerable pyroptosis occurring amongst thymocytes after acute injury caused by TBI (**Fig. 1D**). In fact, direct comparison revealed similar magnitude of activation of both caspase-3 and caspase-1 after damage in DP thymocyte (**Fig. 1E**). Consistent with this induction of immunogenic form of cell death, we found increased release of lactate dehydrogenase in the thymus after TBI (**Fig. 1F**) as well as increased levels of gasdermin D (**Fig. 1G**), all suggesting a preferential induction of pyroptosis following acute damage.

**Figure 1.**
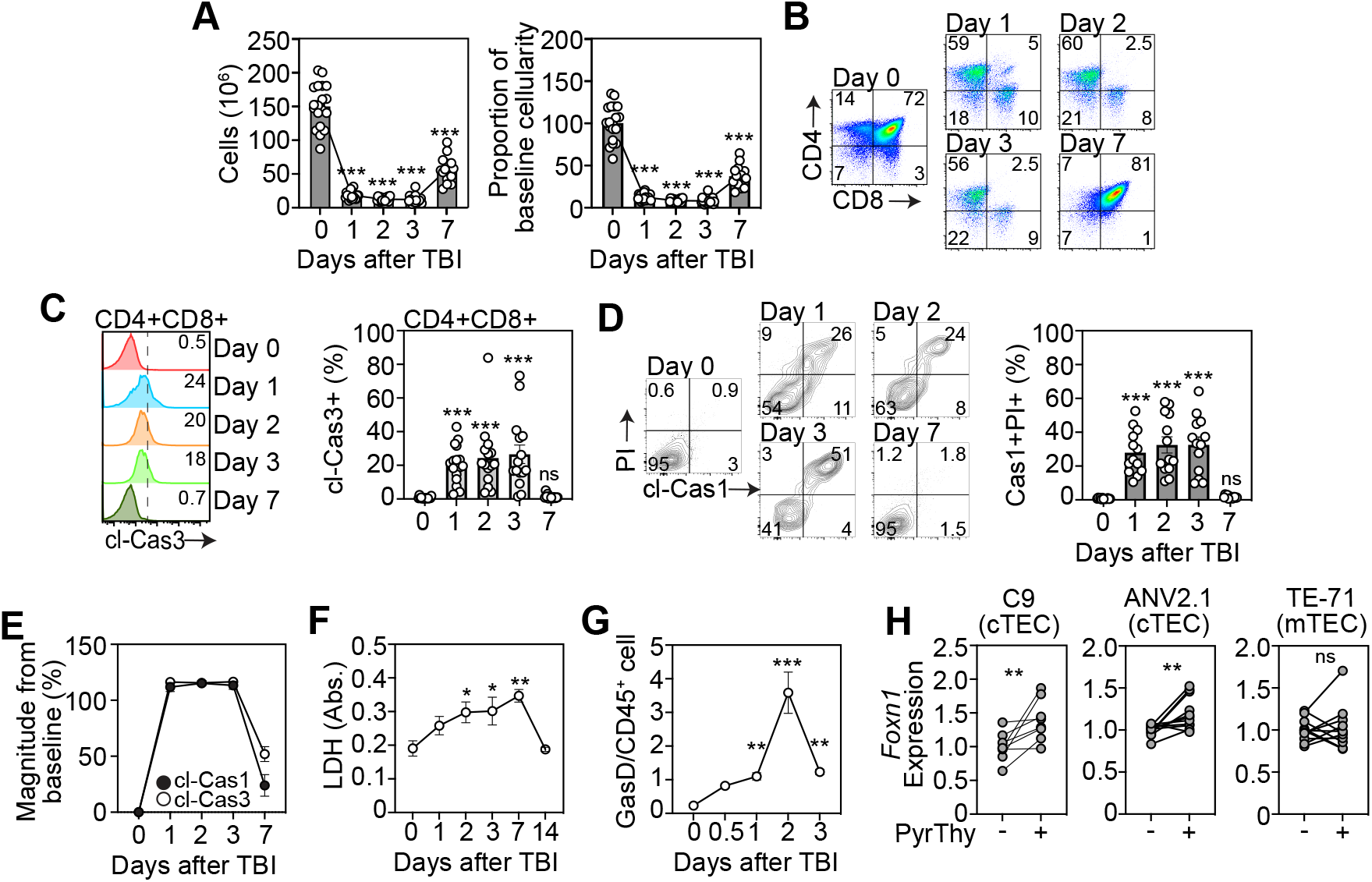
A switch from apoptotic to pyroptotic DP thymocytes triggers thymus regeneration. **A-E,** Thymus was analyzed from 6-8 week old C57/BL/6 mice at days 0,1 2, 3 and 7 following sublethal total body irradiation (TBI, 550cGy). **A**, Total thymic cellularity and proportion of cellularity as a function of baseline cellularity (n=15-19/timepoint from 5 independent experiments). **B**, Concatenated flow cytometry plot of CD4 vs CD8 (gated on viable CD45+ cells) (n=9-13 from 3-4 independent experiments). **C**, Concatenated flow cytometry plot from one experiment showing cleaved caspase-3 on DP thymocytes (Gated on CD45+CD4+CD8+ cells) and bar graph showing proportion of cleaved-cas3+ DP thymocytes (n=15-16/timepoint from 5 independent experiments). **D**, Concatenated flow cytometry plot from one experiment showing cleaved-caspase1 and PI (gated on CD45+CD4+CD8+ cells) and bar graph with proportions (n=13/timepoint from 4 independent experiments). **E**, Magnitude of change in expression of cleaved-caspase-1 and cleaved-caspase-3 in DP thymocytes after TBI. (n=10-15/timepoint/condition from 4-5 independent experiments). **F**, Lactate dehydrogenase levels were measured in the thymus supernatant of mice (n=4 mice/group). **G**, Gasdermin D levels were measured in CD45+ cells from the thymus at days 0,1 2, 3, 7, and 14 post TBI (n=3-4 mice/group from 2-3 independent experiments). **H**; Cells were co-cultured with freshly isolated thymocytes treated to induce or inhibit pyroptosis and Foxn1 expression was measured by qPCR, 20 h after co-culture, in 1C9s (n =8-11 thymuses from 3 separate experiments), ANV42.1 (n=8 thymuses from 3 separate experiments), and TE-71 (n=11 from 3 separate experiments). Data represents mean ± SEM.

Since apoptosis is largely suppressive tissue regeneration in the thymus^10^, we hypothesized that lytic cell death of DPs may be beneficial to regeneration. Moreover, as thymocyte depletion precedes the period of epithelial regeneration largely driven by enhanced *Foxn1* transcription^21^, we tested if pyroptotic thymocytes could directly influence *Foxn1* expression in TECs. To do this we induced pyroptosis in freshly isolated thymocytes *ex vivo* using Nigericin and LPS and co-cultured the dying cells with cortical thymic epithelial cells (cTECs, using the 1C9 and ANV42.1 cell lines) and medullary thymic epithelial cells (mTEC, using the TE-71 cell line) and quantified *Foxn1* expression (**Fig. 1H**). Using this approach, we could demonstrate that the presence of pyroptotic thymocytes directly led to upregulation of *Foxn1* transcription in cTECs but not in mTECs (**Fig. 1H**). This cell-specific regulation of Foxn1 was notable given that we have previously shown that Foxn1 upregulation during endogenous regeneration after damage is largely restricted to cTECs^8^.

### Mitochondrial dysregulation facilitates pyroptosis in DPs

In addition to the critical role of mitochondria in cellular metabolism, the mitochondria is a gatekeeper of cell death and dysregulated mitochondrial bioenergetics can lead to the induction of intrinsic apoptosis or pyroptosis^22, 23, 24, 25^. As thymocytes are undergoing such high levels of homeostatic cell death we sought to understand if metabolic adaptations were steering cell death preferences after damage. First, measuring mitochondrial membrane potential using TMRE revealed a marked induction of mitochondrial membrane hyperpolarization (**Fig. 2A**), correlating to increased cleaved caspase 1 (cl-caspase 1) levels (**Fig. 2B**). Importantly, this enhanced mitochondrial activation was resolved by day 7 following damage, in line with what we observed with caspase 1 cleavage and the re-establishment of apoptosis:pyroptosis balance (**Fig. 1E**). Additional evidence of an acute damage-induced dysregulated metabolic phenotype in DPs was revealed with increased mitochondrial mass in DPs after acute damage, which could also be positively correlated with Cas1 activation (**Fig. 2C-D**).

**Figure 2.**
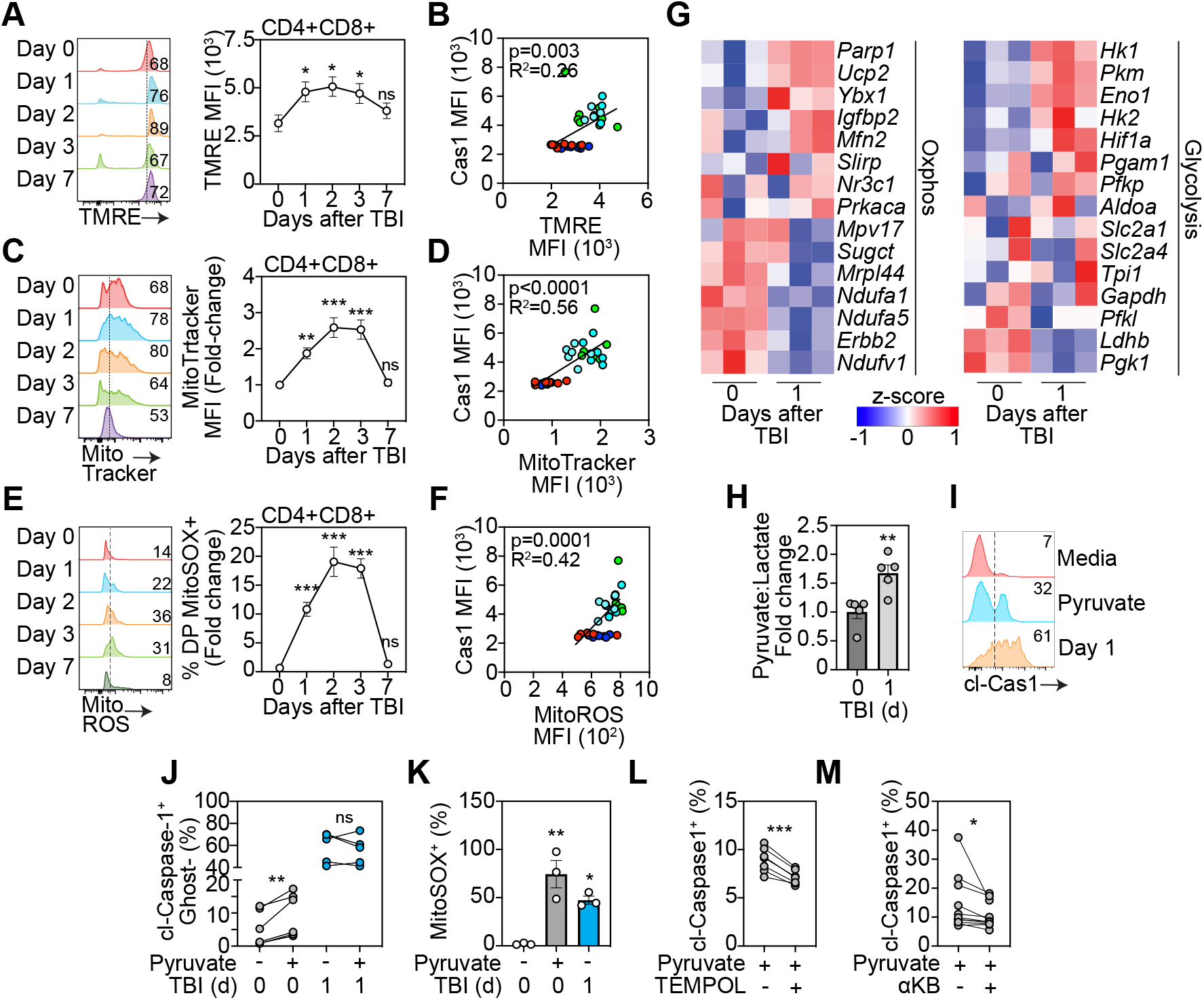
Dysregulated metabolism redirects pyruvate to fuel OXPHOS in thymocytes after damage. **A-F,** Mitochondrial function was analyzed in the thymus isolated from 6-8 week old C57/BL/6 mice at days 0,1 2, 3 and 7 following sublethal total body irradiation (TBI, 550cGy). **A**, Mitochondrial membrane potential assessed by staining of TMRE. Concatenated histogram of TMRE on DP thymocytes (left), quantification of TMRE+ proportions (right) (n=8-12 mice from 3 separate experiments). **B**, Correlation of TMRE expression with Caspase-1 MFI (n=3-8 from 3 independent experiments). **C**, Mitochondrial mass assessed by Mitotracker Green. Concatenated histogram of TMRE on DP thymocytes (left), quantification of TMRE+ proportions (right) (n=9-10 mice from 3 independent experiments). **D**, Correlation of TMRE expression with Caspase-1 MFI (n=6/timepoint from 3 independent experiments). **E**, Mitochondrial ROS was assessed by staining for MitoSOX. Concatenated histogram of TMRE on DP thymocytes (left), quantification of TMRE+ proportions (right) (n=5-7 mice from 2 separate experiments). **F**, Correlation of TMRE expression with Caspase-1 MFI (n=5-7/timepoint from 2 independent experiments). **G**, RNA seq was carried out on FACS purified CD4+CD8+ thymocytes from untreated and TBI-treated (1 day post TBI) mice (n=3/group). Displayed are heatmaps for expression of key genes involved with OXPHOS and glycolysis. **H**, Intracellular lactate and puyruvate levels were measured in freshly isolated thymocytes from untreated and TBI-treated mice (n=5 mice/group from 2 separate experiments). **I-K**, Thymocytes isolated from mice at days 0 or 1 post TBI were incubated with RPMI supplemented with pyruvate (5 mM) for 4 h. **I**, Concatenated histogram showing expression of cleaved-caspase-1 on DP thymocytes in cells incubated with meadia alone, high pyruvate (on day 0 thymocytes) or normal media with thymocytes isolated at d1 following TBI. **J**, Proportion of Cleaved-caspase-1+GhostDye+ cells (n=6 mice from 2 separate experiments). **K**, Proportion of mitoSOX+ DP thymocytes (n=6 mice from 2 separate experiments). **L**, Freshly isolated thymocytes from untreated mice were incubated with RPMI supplemented with pyruvate (5 mM) for 4 h plus TEMPOL (100 µM). cl-caspase 1 levels were measured using flow cytometry (n=6 mice from 2 separate experiments). **M**, Freshly isolated thymocytes from untreated mice were incubated with RPMI supplemented with pyruvate (5 mM) for 4 h plus α-ketobutyrate (200 µM). cl-caspase 1 levels were measured using flow cytometry (n=13-14 mice from 4 separate experiments). Data represents mean ± SEM.

### Increased mitochondrial ROS triggers pyroptosis in thymocytes

Bidirectional communication between the mitochondria and the NLRP3 inflammasome has been well characterized and can induce activation of NLRP3 signaling^26, 27, 28^, while concurrently facilitating a lack of mitophagy driven by cleavage of caspase 1^29, 30^. This positive feedback loop perpetuates the accumulation of ROS-producing dysfunctional mitochondria due to a lack of mitophagy, which in turns continues to initiate NLRP3-induced pyroptotic cell death^25, 31^. Next, we hypothesized that transiently enhanced mitochondrial activation in DPs led to increased production of mitochondrial ROS (mitoROS) providing a trigger for pyroptosis. Consistent with mitochondrial hyperpolarization, mitochondrial mass and cl-caspase 1 levels, there was a transient and precipitous elevation in mitoROS levels after damage, again correlating with caspase-1 cleavage (**Fig. 2E-F**). This increase in mitoROS was coupled with a robust and acute induction in glutathione levels (**Fig. S1**), suggesting antioxidant pathways are upregulated. RNA seq analysis of DP thymocytes at baseline and 24 hours after TBI revealed an increase in *Ucp2* and *Mitofusin 2* (**Fig 2G**), which are central facilitators of proton leakage and Nlrp3 activation^32, 33^. RNA sequencing on DP thymocytes also revealed an enrichment for genes regulating OXPHOS (*Igf2bp2, Ybx1, Ucp2*) and, importantly, downregulation of pyruvate processing to lactate (*Ldhb*), pointing to a redirection of pyruvate to fuel OXPHOS (**Fig. 2G**). Of note, genes encoding key glycolysis enzymes, such as *Hk1* and *Hk2*, were upregulated after TBI suggesting an increase in glucose uptake, providing increased levels of pyruvate as fuel for mitochondrial metabolism^34, 35^.

In order to demonstrate that dysregulated metabolism was driving this shift from apoptosis to pyroptosis after damage, we examined the role of increased mitochondrial metabolism on cell death in thymocytes. Firstly, to assess any damage-induced alterations in glycolytic flux we measured pyruvate and lactate levels in thymocytes at rest and 24 h after damage and the ratio of pyruvate to lactate was significantly increased early after damage (**Fig. 2H**); strengthening our findings of a damage-induced metabolic shift away from glycolysis and towards OXPHOS. Next, to determine if enhancing mitochondrial respiration by increasing pyruvate could induce caspase 1 cleavage and cell death, we incubated freshly isolated thymocytes *ex vivo* with high levels of pyruvate (5 mM). This approach demonstrated that pyruvate induced caspase 1 cleavage in thymocytes, but this could not be induced in thymocytes isolated from mice given TBI (**Fig. 2I-J**), suggesting a zenith of pyroptosis, possibly due to the saturation of pyruvate and mitochondrial activity acutely after damage. Consistent with this proposed mechanism, treatment of thymocytes with high levels of pyruvate strongly induced mitoROS (**Fig. 2K**) and targeting mitoROS with the inhibitor TEMPOL reduced cl-caspase 1 levels in thymocytes under pyruvate pressure (**Fig. 2L**). Finally, blocking pyruvate conversion to acetyl co-A with α-ketobutyrate (α-KB) reduced cl-caspase 1 levels, demonstrating a role of the TCA cycle in pyroptosis induction in thymocytes (**Fig. 2M**). These findings were consistent with a demonstration that fueling increased proton leakage and increased mitoROS triggers pyroptosis in thymocytes after damage, confirming DPs preferentially undergo pyroptotic cell death after damage facilitated by increased pyruvate-induced production of mitoROS.

### Extracellular ATP induces Foxn1 expression in cTECs

Pyroptotic cell death produces a plethora of molecules that act as ligands and messengers to facilitate communication with neighboring cells^36,37,38^. We have previously shown that extracellular Zn^2+^ can act as a damage-associated molecular pattern (DAMP) after acute damage, inducing expression of the pro-regenerative molecule BMP4 in endothelial cells via the receptor GPR39^39^.

However, activation of GPR39 on TECs failed to induce *Foxn1* expression. We thus sought to identify specific DAMPs that could trigger the induction of *Foxn1* transcription, specifically focusing on cTECs. To this end, we tested the response of the cTEC cell line (1C9) to a panel of DAMPs and identified ATP to be a strong inducer of *Foxn1* transcription (**Fig. 3A**). This finding was confirmed in another cTEC cell line (ANV42.1) (**Fig. 3B**) and, importantly, freshly isolated human TECs (**Fig. 3C**).

**Figure 3.**
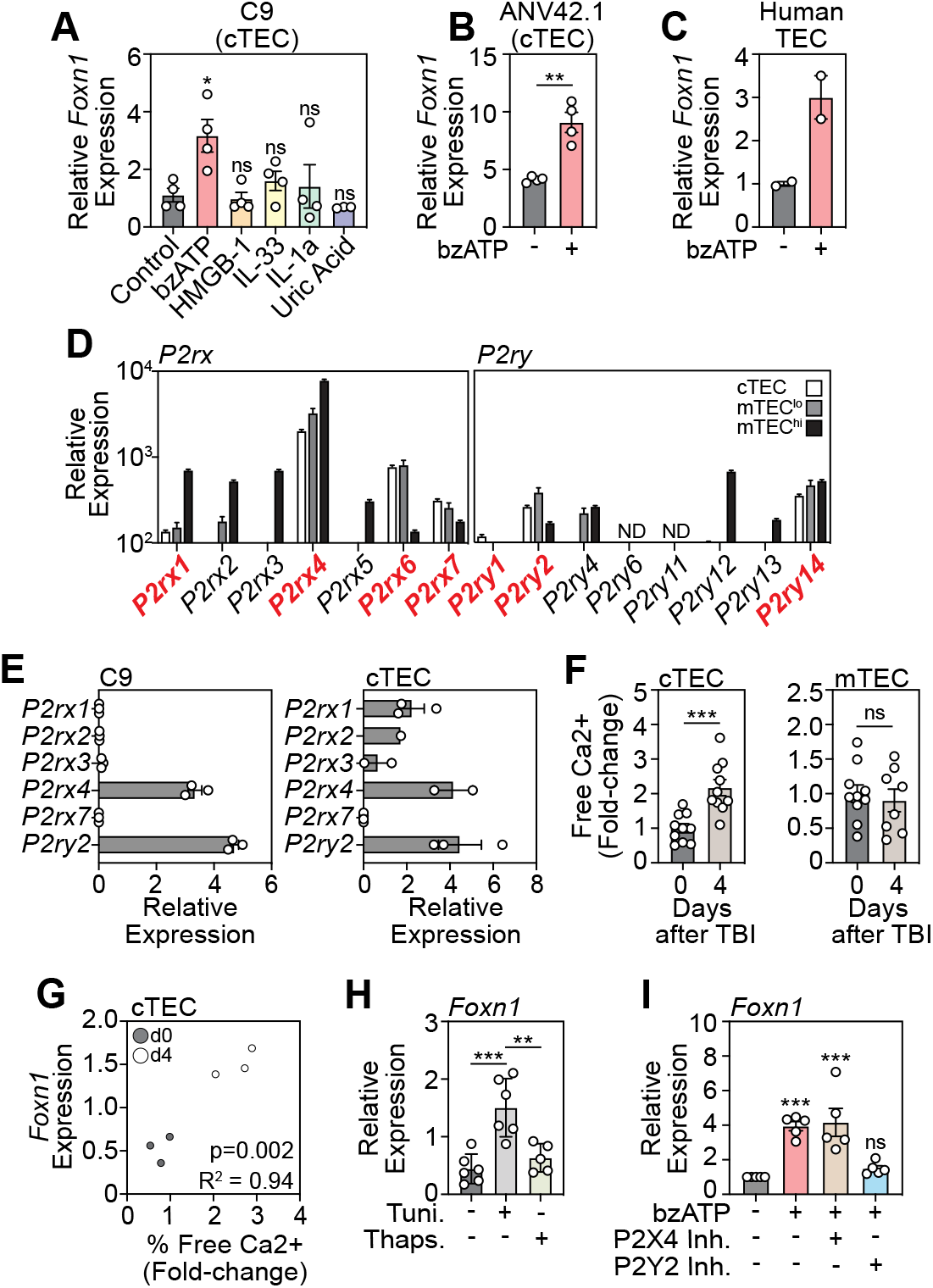
Activation of the P2Y2 receptor with extracellular ATP induces FOXN1 in cTECs. **A,** The cTEC cells line 1C9 was treated with a panel of DAMPs and *Foxn1* transcription was measured in by qPCR 20 h following incubation (n=3-4 separate experiments). **B**, A second cTEC cell line (ANV42.1) was treated with ATP (100 µM) and *Foxn1* expression was measured after 20 h (n=3 separate experiments). **C**, Freshly isolated human cTECs were treated with ATP (100 µM) and *Foxn1* expression was measured after 20 h (n= 2). **D**, cTECs, MHCII^hi^ mTEC, and MHCII^lo^ mTEC were isolated from untreated 6wo C57BL/6 mice and RNA sequencing performed. Displayed is expression of purinergic receptor family members (n=3/cell population). **E**, Purinergic receptor expression in FACS purified cTECs from untreated mice measured by qPCR (n=2-3 pooled mouse thymuses). **F**, Intracellular free Ca^2+^ levels were measured by flow cytometry in untreated and TBI-treated (day 4 post TBI) cTECs (n=10 mice from 2 separate experiments) and mTECs (n=8-10 mice from 2 separate experiments). **G**, Correlation of free Ca^2+^ and Foxn1 expression at days 0 and 4 after SL-TBI (n=3/timepoint). **H**, 1C9 (cTEC) were treated with tunicamycin (1 µM) or thapsigargan (100 nM) for 20 h and *Foxn1* expression was measured by qPCR (n=4 independent experiments). **I**, 1C9 (cTECs) were treated with ATP and either antagonists for P2Y2 or P2X4 and *Foxn1* expression was measured by qPCR 20 h after incubation (n=5 separate experiments). Data represents mean ± SEM.

ATP is a ligand for cell surface purinergic receptors and can activate downstream signaling pathways that either induce the influx of extracellular Ca^2+^ or promote the efflux of ER Ca^2+^ via G-coupled signaling^40, 41, 42^. Previous studies have found that purinergic receptor expression is heterogeneous between thymic epithelial cell subsets, with widespread expression of both P2Y and P2X receptors expressed among all subsets of TECs^43^. Consistent with this, specific analysis of TEC subsets by RNA sequencing revealed expression of multiple P2X and P2Y receptors across cTECs and mTECs, with expression on cTECs limited to *P2rx1, P2rx4, P2rx6, P2rx7, P2ry1, P2ry2*, and *P2ry14* (**Fig. 3D**). Baseline expression levels of purinergic receptors were confirmed by qPCR on the 1C9 (cTEC cell line), and on freshly isolated murine cTECs (**Fig. 3E**). Next, as P2 receptor activation induces a downstream increase in intracellular Ca^2+^ levels, we measured Ca^2+^ levels in cTECs and mTECs after damage and demonstrated an increase in Ca^2+^ in cTECs but not mTECs (**Fig. 3F**), with positive correlation between Ca^2+^ levels with *Foxn1* expression (**Fig. 3G**). This data is consistent with the cell-specific effects of pyroptotic thymocytes on *Foxn1* expression in cTECs, suggesting cTECs are central gatekeepers of the ATP-mediated regenerative response. As P2X and P2Y receptor activation regulate Ca^2+^ levels differently^40^, we assessed the effect of ATP on both Ca^2+^ influx and efflux within cTECs. To do this we treated cTECs with tunicamycin, to induce ER release of Ca^2+^ into the cytosol, or thapsigargan, to inhibit ER Ca^2+^ release, and revealed that flooding the cell with Ca^2+^ led to enhanced *Foxn1* expression, while attenuating Ca^2+^ levels restored *Foxn1* expression to baseline (**Fig. 3H**). These results strongly suggested the role of P2Y receptors in mediating the FOXN1 promoting effects of extracellular ATP, however, we further sought to refine our target and identify which P2 receptor was critical to mediate this effect. To confirm this, we treated cTECs with ATP in the presence of antagonists for P2Y2, and P2X4, as P2X4 is highly expressed on cTECs although does not induce Ca^2+^ efflux and demonstrated that inhibition of P2Y2 attenuated ATP-mediated *Foxn1* induction, mirroring the effects ATP elicits as an extracellular DAMP after acute insult (**Fig. 3I**).

### Specific activation of P2Y2 receptors can enhance Foxn1 expression and boost thymic function after acute injury

P2 antagonists are of increasing interest therapeutically, with a focus on developing analogous molecules to inhibit or promote these druggable targets in many disease settings, such as epilepsy^44^, rheumatoid arthritis^45^ and ischemic cardiac injury^46^. Moreover, clinical trials have been carried out using antagonists for P2X3^47^, P2X7^48, 49^ and P2Y12^50^. To test if P2Y2 could be targeted to enhance FOXN1 we obtained a specific P2Y2 agonist and further demonstrated that stimulation of cTECs with a P2Y2 agonist induced *Foxn1* expression, and inhibition of P2Y2 ablated this response (**Fig. 4A**). We sought to translate our molecular target discovery findings into a therapeutic strategy *in vivo* to test if P2Y2 agonism could enhance thymic regeneration. To do this we treated C57BL/6 mice with SL-TBI and administered the P2Y2 agonist UTPγS intraperitoneally at day 1 following damage and assessed thymic cellularity 13 days after damage and confirmed that UTPγS could enhance thymus regeneration after acute damage (**Fig. 4B**). Additionally, *in vivo* treatment with UTPγS had a global impact on thymocyte populations, with increased regeneration of DPs, CD4+ and CD8+ thymocytes and superior regeneration of the TEC compartment (**Fig. 4C)**.

**Figure 4.**
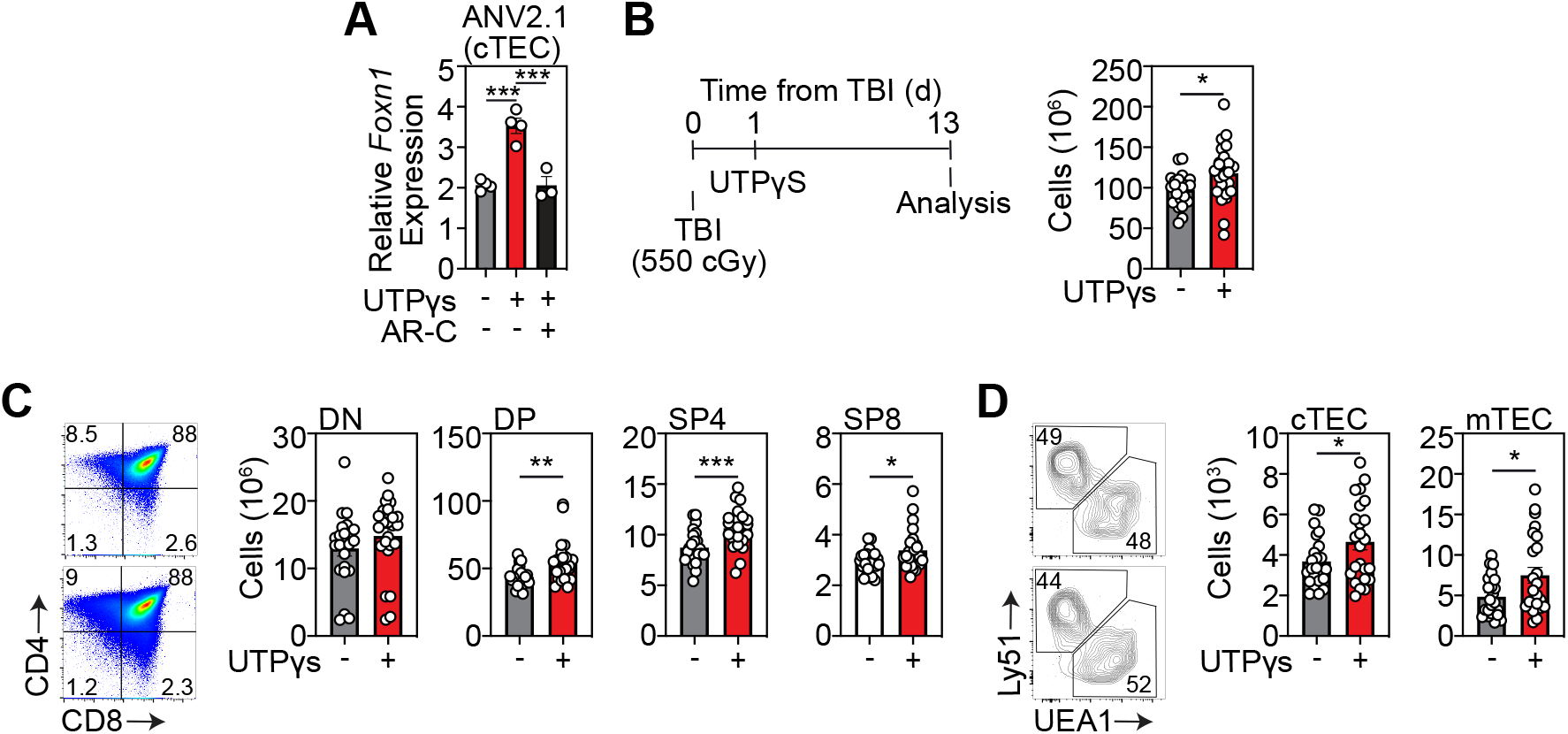
Activation of the P2Y2 enhances thymus regeneration after damage. **A,** ANV42.1 (cTEC) cells were treated with the P2Y2 agonist UTPγS and the P2Y2 antagonist ARC-118925XX for 20 h and *Foxn1* expression was measured by qPCR after 20 h (n=3 separate experiments); **B-D**, 6wo C57BL/6 mice were treated with UTPyS (1 mg/kg) IP at day 1 following SL-TBI and the thymuses were harvested at day 13. **B**, Total thymus cellularity. **C**, Number of CD4-CD8-double negative (DN), DP, of CD4 or CD8+ single positive (SP4 or SP8, respectively) thymocytes (n=23-24 mice from 3 independent experiments). Data represents mean ± SEM.

## DISCUSSION

Endogenous thymic regeneration engages complex multicellular signaling networks that intricately communicate within cellular niches to repair and replenish peripheral T cell reconstitution. Here, we demonstrated that an induction of pyroptosis as a preferred cell death mechanism provides the critical ligand ATP, that stimulated P2Y2 receptors on cTECs to promote FOXN1 transcription and enhance regeneration of the thymus. Moreover, we uncovered a cell specific mechanism of metabolically regulated thymus regeneration that is centered on streering pyruvate processing to induce lytic cell death by dysregulating mitochondrial metabolism, acutely increasing mitoROS and triggering NLRP3 activation and pyroptosis. Here we identified the effects of acute damage on the thymocyte metabolic landscape and cell fate.

The thymus is highly hypoxic^51, 52^, and thymocytes undergo dynamic alterations in respiration during development (specifically between DN and DP stages)^53^, pointing to their metabolic plasticity. Here we identified the effects of acute damage on the metabolic landscape of thymocytes and revealed a that increased levels of pyruvate are redirected toward mitochondrial respiration, reducing glycolysis. Disruption of glycolytic flux has been shown to trigger pyroptosis^54^, and our data demonstrates that an acutely altered metabolic profile in DP thymocytes drives pyroptosis in the thymus and is rapidly resolved as regeneration is initiated. Accompanying damage-induced acute hyperpolarization of the mitochondrial membrane, increased mitochondrial mass and mitochondrial ROS in DP thymocytes, our RNA sequencing data showed an upregulation of genes encoding UCP2 and PARP1, key negative regulators of oxidative stress^55, 56^, and in Mitofusin 2 which governs mitochondrial integrity^57^. Importantly, our gene signature of damage in DP thymocytes revealed that several genes encoding enzymes critical for glucose processing to pyruvate, such as HK1, HK2, PKM and HIF1α were upregulated concurrently with a downregulation in genes regulating pyruvate conversion to lactate (*Ldhb*). Functionally resulting in a higher pyruvate to lactate ratio, and redirection towards mitochondrial respiration. ROS reacts with the NLRP3 inflammasome and drives pyroptotic cell death^25, 58, 59, 60, 61, 62^. Here we confirmed this in the thymus and demonstrated that pyruvate drives this increase in mitochondrial ROS that further triggers caspase-1 cleavage and pyroptosis, strengthening the case for a central role in redirection away from glycolysis as a trigger from thymocyte cell death.

Both intracellular and extracellular ATP has been previously identified to play a role in tissue repair^63, 64, 65^, and importantly purinergic receptors are identified to mediate the extracellular ATP response^66^, with interest in pharmacologically targeting these receptors to enhance wound repair^67, 68^. Activation of purinergic receptors mobilizes intracellular Ca^2+^ in epithelial cells^69^. Here we identified that inhibiting Ca^2+^ efflux from the ER, using thapsigargan, downstream of ATP treatment prevented FOXN1 transcription, which led us to further assess P2Y2 as a specific target. Moreover, P2Y2 has been identified to mediate migration and repair of epithelial cells^70^. Purinergic receptors have been trialed for a range of diseases, for example P2X7 antagonists are being tested is a Phase 2a clinical trial to treat Crohns disease, while targeting P2X7 to treat rheumatoid arthritis failed to show significance in phase 2 clinical trial^71^, and the P2Y2 agonist Diquafosol is currently being tested for the treatment of dry eye^72^. We identified that P2Y2 agonism promotes FOXN1 transcription specifically in cTECs and that competition with an antagonist quenches this effect, pointing to receptor specificity. Moreover, our pre-clinical data demonstrates that the P2Y2 agonist UTPγS promotes superior regeneration in acutely injured mice, promoting recovery of thymocyte and both cTEC and mTEC compartments, which is vital for continued maintenance and functioning of the thymus.

While much remains to be understood regarding mitochondrial regulation of cell death and differentiation in models of chronic damage such as age, these data underline an important mechanism of recovery from acute damage that highlights the significance of metabolic governance of immune function. The question of other fuel sources that drive mitochondrial dysfunction during acute damage, such as lipids or glutamine, is outstanding and may potentially reveal disease specific damage-responses of the thymus, specifically as lipid metabolism has been identified to play a central role in hematopoiesis and T cell differentiation^73^. However, as these metabolic phenotypes are likely to be variable between cellular compartments, and with our data clearly demonstrating a central role of pyruvate in mitochondrial induced pyroptosis, the convergence of these pathways on mitochondrial ROS is central to pyroptotic driven regeneration. In conclusion, these data describe a complex molecular architecture that govern thymus regeneration and not only provides a platform for therapeutic target discovery and intervention towards enhancing immune function, but also contributes to regenerative medicine by unravelling novel mechanisms of metabolically regulated endogenous tissue regeneration which may be applicable across multiple tissues.

## MATERIALS AND METHODS

### Mice

Inbred male and female C57BL/6J mice were obtained from the Jackson Laboratories (Bar Harbor, USA) and all experimental mice were used between 6-8 weeks old. To induce thymic damage, mice were given sub-lethal total body irradiation (SL-TBI) at a dose of 550 cGy from a cesium source mouse irradiator (Mark I series 30JL Shepherd irradiator) with no hematopoietic rescue. Mice were maintained at the Fred Hutchinson Cancer Research Center (Seattle, WA), and acclimatized for at least 2 days before experimentation, which was performed per Institutional Animal Care and Use Committee guidelines.

### Reagents

Cells were stained with the following antibodies for analysis CD3-FITC (35-0031, Tonbo Bioscience), CD8-BV711 (100748, BioLegend), CD4-BV650 (100546, BioLegend), CD45-BUV395 (565967, BD Biosciences), CD90-BV785 (105331, BioLegend), MHC-II-Pac Blue (107620, BioLegend), EpCAM-PercPe710 (46-5791-82, eBioscience), Ly51-PE (12-5891-83, eBioscience), UEA1-FITC (FL-1061, Vector Laboratories), TCRbeta-PECy7 (109222, BioLegend), CD62L-APC-Cy7 (104427, BioLegend), CD44-Alexa Fluor RTM700 (56-0441-82, BioLegend), CD25-PercP-Cy5.5 (102030, BioLegend). Flow cytometry analysis was performed on an LSRFortessa X50 (BD Biosciences) and cells were sorted on an Aria II (BD Biosciences) using FACSDiva (BD Biosciences) or FlowJo (Treestar Software).

### Thymus digestion and cell isolation

Single cell suspensions of freshly dissected thymuses were obtained and either mechanically suspended or enzymatically digested as previously described^5, 74^ and counted using the Z2 Coulter Particle and Size Analyzer (Beckman Coulter, USA). For studies sorting rare populations of cells in the thymus, multiple identically treated thymuses were pooled so that sufficient number of cells could be isolated; however, in this instance separate pools of cells were established to maintain individual samples as biological replicates.

### Cell death assays

Thymuses from untreated and SL-TBI treated mice were harvest, enzymatically digested and stained with cell surface markers for thymus populations. Cells were further stained for caspase 1 cleavage with Caspase-1 (active) Staining Kit (Abcam, ab219935), or fixed for caspase 3 analysis using Cleaved Caspase-3 (Asp175) Antibody (Alexa Fluor® 488 Conjugate) (Cell signaling, 9669S). Apoptosis and pyroptosis was assessed by adding Annexin V-FITC (Biolegend, 640906), Annexin V binding buffer (BioLegend, 422201) and Propidium Iodide (Invitrogen, BMS500PI). Gasdermin D was measured in freshly isolated thymocytes using Gasdermin D (mouse) ELISA Kit (Adipogen Life Sciences, AG-45B-0011-KI01). Lactate dehydrogenase was assessed from the supernatant of harvested thymocytes using Lactate Dehydrogenase assay (Abcam, ab102526).

### In vitro assays

*Co-culture assays:* thymocytes were isolated from untreated C57BL/6 mice and incubated with Nigericin (10 µM, Tocris, 4312) and LPS (1 ng/ml, Invivogen, tlrl-eblps) for 3 h and co-cultured with 1C9s, ANV42.1 or TE-71 cell lines for 20 h before lysis for qPCR. *DAMP stimulation:* 1C9s were stimulated with ATP (100 µM, Tocris 3312), HMGB1 (1 µg/ml, Abcam, ab78940), IL1a (50 ng/ml,Tocris, 400-ML-005/CF), IL-33 (50 ng/ml, Tocris, 3626-ML-010/CF), or uric acid (50 µg/ml, Sigma, U2625) for 20 h and lysed for qPCR analysis. *UTPγS assays:* 1C9s or ANV42, cells were stimulated with UTPγS (100 µM, R&D Systems, 3279), or UTPγS plus AR-C 118925XX (20 µM, Tocris, 4890) for 20 h before lysis for qPCR. BzATP triethylammonium salt (100-300 µM, Tocris, 3312), was used for ATP stimulation.

### qPCR and RNA sequencing

RNA was extracted from exECs or DCs using a RNeasy Mini kit (74104, Qiagen), and from sorted cells using a RNeasy Plus Micro kit (74034, Qiagen). cDNA was synthesized using the iScript gDNA Clear cDNA Synthesis kit (1725035, Bio-Rad, USA) and a Bio-Rad C1000 Touch ThermoCycler (Bio-Rad). RNA expression was assessed in the Bio-Rad CFX96 Real Time System (Bio-Rad), using iTaq Universal SYBR Green Supermix (1725122, Bio-Rad), and the following primers: β-Actin (F 5’-CACTGTCGAGTCGCGTCC-3’; R 5’-TCATCCATGGCGAACTGGTG-3’); PrimePCR™ SYBR® Green Assay: Foxn1, Mouse (Biorad, 10025637, qMmuCED0044924).

RNAseq was performed on freshly isolated and FACS purified CD4+CD8+ thymocytes. To obtain sufficient RNA for every timepoint, thymi of 2 mice were pooled for untreated mice and 6 for irradiated mice. All samples underwent a quality control on a bioanalyzer to exclude degradation of RNA.

### Ex vivo metabolic assays

Thymuses from untreated and SL-TBI treated mice were harvested and enzymatically digested and stained from flow cytometry analysis of thymocyte populations as above. Further analysis of mitochondrial bioenergetics were assessed using TMRE (Abcam, ab113852), MitoTracker™ Green FM (Invitrogen, M7514), MitoSOX™ Red Mitochondrial Superoxide Indicator (ThermoFisher, M36008), and Intracellular glutathione (GSH) Detection Assay Kit (Abcam, ab112132). Thymocytes were isolated from untreated and TBI-treated mice and intracellular pyruvate and lactate levels were measured by absorbance using Pyruvate Assay kit (Abcam, ab65342) or Lactate-Glo™ Assay (Promega, J5021). Thymocytes were isolated from untreated and TBI-treated mice and incubated in RPMI with 5 mM sodium pyruvate (Gibco, 11360070) for 3 h at 37 °C and stained for flow cytometry analysis and cl-caspase 1 levels. Cells were further incubated with 5 mM sodium pyruvate plus 200 µM α-ketobutyrate (Sigma-Aldrich) or 100 µM TEMPOL (Tocris, 3082) for 3 h and cells were prepared for flow cytometry analysis.

### Intracellular Ca2+ assay

The thymuses from untreated and SL-TBI-treated mice were harvested and processed for flow cytometry as above. The intracellular Ca^2+^ dye BAPTA-AM/Indo-AM was added (Sigma-Aldrich). Unbound intracellular Ca^2+^ was assessed in cTECs and mTECs by measuring BAPTA-AM levels on the BUV-496 filter.

### In vivo UTPγS administration

For *in vivo* studies of UTPγS administration, mice were given SL-TBI (550cGy) and subsequently received intraperitoneal injections of 1 mg/kg UTPγS (R&D systems, 3279), or 1x PBS as control, on day 1 following TBI. Thymuses were harvested 13 days after SL-TBI and cellularity was assessed and populations were analyzed by flow cytometry.

### Statistics

All analysis between two groups was performed with a non-parametric Mann-Whitney test. Statistical comparison between 3 or more groups in Figs. 1A, 1C, 1D, 1E, 1F, 1G, 2A, 2B, 2C, 2E, 3A, 3H, 3I, and 4A were performed using a one-way ANOVA with Tukey’s multiple comparison post-hoc test. All statistics were calculated using Graphpad Prism and display graphs were generated in Graphpad Prism or R.

## ACKNOWLEDGEMENTS

We gratefully acknowledge the assistance of the Flow Cytometry and Comparative Medicine Core Facilities; and the support of the Immunotherapy Integrated Research Center at the Fred Hutchinson Cancer Research Center. We are grateful to Fionnuala Morrish and Carla Jaeger for discussions regarding metabolic assays. This research was supported by National Institutes of Health award numbers R00-CA176376 (J.A.D.), R01-HL145276 (J.A.D.), R01-HL165673 (J.A.D.), U01-AI70035 (J.A.D.), Project 2 of P01-AG052359 (J.A.D.), and the NCI Cancer Center Support Grant P30-CA015704. Support was also received from a Scholar Award from the American Society of Hematology (J.A.D.); the Mechtild Harf (John Hansen) Award from the DKMS Foundation for Giving Life (J.A.D.); the Cuyamaca Foundation (J.A.D.), and the Bezos Family Foundation (J.A.D.). S.K. was supported by a New Investigator Award from the American Society for Transplantation and Cellular Therapy and Pilot Funding from the Cooperative Center for Excellence in Hematology (Fred Hutchinson Cancer Research Center) award number U54 DK106829.

## AUTHOR CONTRIBUTIONS

S.K. and J.A.D. conceived of the idea of this manuscript. J.A.D., and S.K. designed, analyzed and interpreted experiments, and drafted the manuscript; C.A.E., K.C., L.I., P.d.R, A.C., K.S.H., C.W.S., and D.G. performed experiments; L.S., E.V., and J.A.D. supervised experiments. All authors contributed to the article and approved the submitted version.

---

**Supplementary Figure 1.**
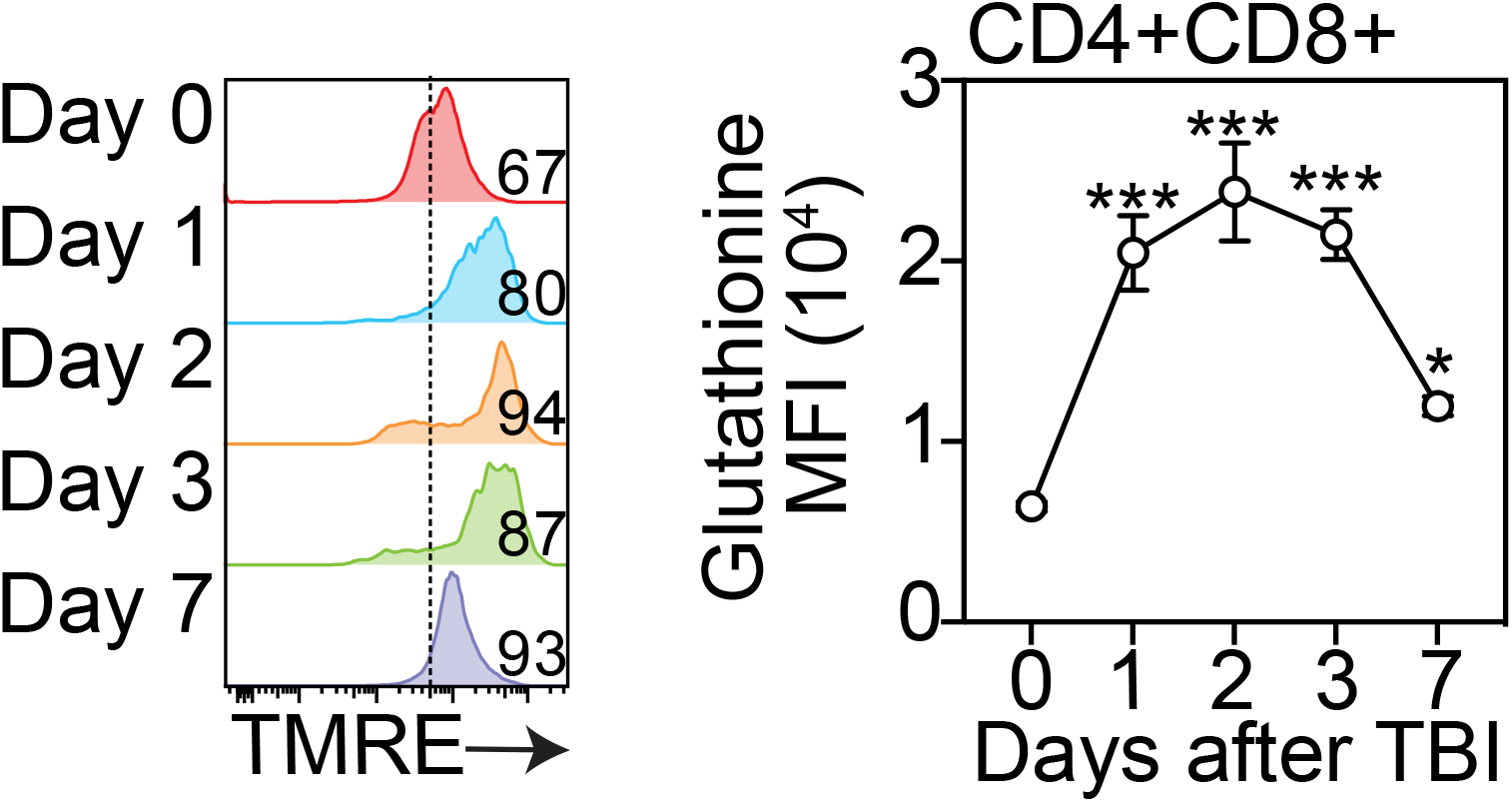
Thymuses from 6-8 week old C57BL/6 mice were harvested at days 0, 1, 2, 3 and 7 following TBI and glutathione levels were measured by flow cytometry in DP thymocytes (n=3 mice).

